# Isolate-anchored comparisons reveal evolutionary and functional differentiation across SAR86 marine bacteria

**DOI:** 10.1101/2024.03.17.584874

**Authors:** Oscar Ramfelt, Kelle C. Freel, Sarah J. Tucker, Olivia D. Nigro, Michael S. Rappé

## Abstract

SAR86 is one of the most abundant groups of bacteria in the global surface ocean. However, since its discovery over 30 years ago, it has remained recalcitrant to isolation and many details regarding this group are still unknown. Here we report the cellular characteristics from the first SAR86 isolate brought into culture, *Candidatus* Magnimaribacter mokuoloeensis strain HIMB1674, and use its closed genome in concert with over 700 environmental genomes to assess the phylogenomic and functional characteristics of this order-level lineage of marine Gammaproteobacteria. The Magnimaribacterales invest significant genomic resources into the capacity for β-oxidation, which is present in most genomes in high gene copy numbers. This cyclical set of reactions is fed by components of cell membranes that includes lipids such as phosphatidylcholine, phosphatidylethanolamine, glycolipids, and sulfolipids. In addition to the widespread capacity to degrade the side chain of steroidal compounds via β-oxidation, several SAR86 sublineages also appear able to fully degrade the steroid polycyclic ring structure as well as other aromatic, polycyclic, and heterocyclic molecules. Read recruitment from publicly available metagenomes reveals that the Magnimaribacterales compose up to 6% of the global surface ocean microbial community. Only a subset of genera drive these high relative abundances, with some more globally dominant and others restricted to specific oceanic regions. *Candidatus* Magnimaribacter mokuoloeensis provides an unprecedented foundation through which to understand this highly abundant yet poorly understood lineage of marine bacteria, and charts a path to bring more representatives of this order into laboratory culture.

## Main

The marine bacterial lineage known as SAR86 was initially discovered over 30 years ago through the application of 16S rRNA gene sequencing to identify microorganisms within natural communities of marine plankton^1–3^; “SAR” for its discovery in a surface seawater sample from the Sargasso Sea, and “86” for clone 86. Soon after its discovery within the open-ocean gyres of the Atlantic and Pacific Oceans^1–3^, diverse relatives of the original gene clone were recovered from around the global ocean, including coastal systems^4–9^. Reports on the abundance of SAR86 by counting cells via fluorescence *in situ* hybridization, quantitative PCR, and other molecular approaches routinely support the view that it is one of the most abundant bacteria in seawater, particularly in the surface ocean^10–16^.

In a landmark study, one of the first applications of environmental genomics revealed a gene encoding the photoactive transmembrane protein proteorhodopsin in a SAR86 genome fragment^17^. While research on proteorhodopsin has flourished^18–21^, a comprehensive picture of SAR86 marine bacteria has been slower to develop. In this study, we provide the first characterization of SAR86 cells in culture and leverage the closed genome sequence of the isolated SAR86 strain in a set of analyses designed to elucidate the evolutionary and functional characteristics of SAR86 bacteria. Despite the ubiquity of SAR86 cells in the global ocean, there is currently little to link extensive 16S rRNA gene-based studies from the past three decades with the genomic diversity uncovered over the past two decades^12,22–25^, and thus no existing backbone from which to interpret relationships between phylogenetic diversity, environmental distribution, and the distribution of functional traits encoded by genomes. Thus, included in our goals were to carefully investigate the HIMB1674 genome along with hundreds of publicly available SAR86 environmental genomes in order to evaluate the evolutionary origins of SAR86 within the *Gammaproteobacteria*, as well as evolutionary relationships between the strain genome and other SAR86 environmental genomes. We also sought to identify the genomic features that typify HIMB1674 and other SAR86 bacteria, as well as features that may distinguish major sublineages within SAR86. Finally, by recruiting sequence reads from publicly available metagenomes from marine plankton, we aimed to provide the first comprehensive analysis unifying evolutionary history with the environmental distribution of SAR86 genotypes.

## Results

### Strain HIMB1674 contains small cells and a genome typical of oligotrophic marine bacteria

A large dilution-to-extinction cultivation experiment using coastal Hawaiʻi surface ocean seawater resulted in 339 isolates that were then identified via short-read 16S rRNA gene sequencing. The affiliation of one strain (HIMB1674) with the SAR86 clade was confirmed by full length 16S rRNA gene sequencing (Fig. 1a). Whole genome sequencing yielded a single closed scaffold of 1.62 Mbp in length and a GC content of 36.4%, containing 1,699 total genes (1,645 protein-coding), one rRNA operon, and a coding density of 96.3% (Table 1). Scanning and transmission electron micrographs consistently revealed strain HIMB1674 to be uniformly small spheroid bacilli (0.46±0.03 × 0.66±0.11 μm; with an estimated biovolume of 0.07±0.01 μm^3^) (Fig. 2a). We have named strain HIMB1674 *Candidatus* Magnimaribacter mokuoloeensis.

**Fig. 1:**
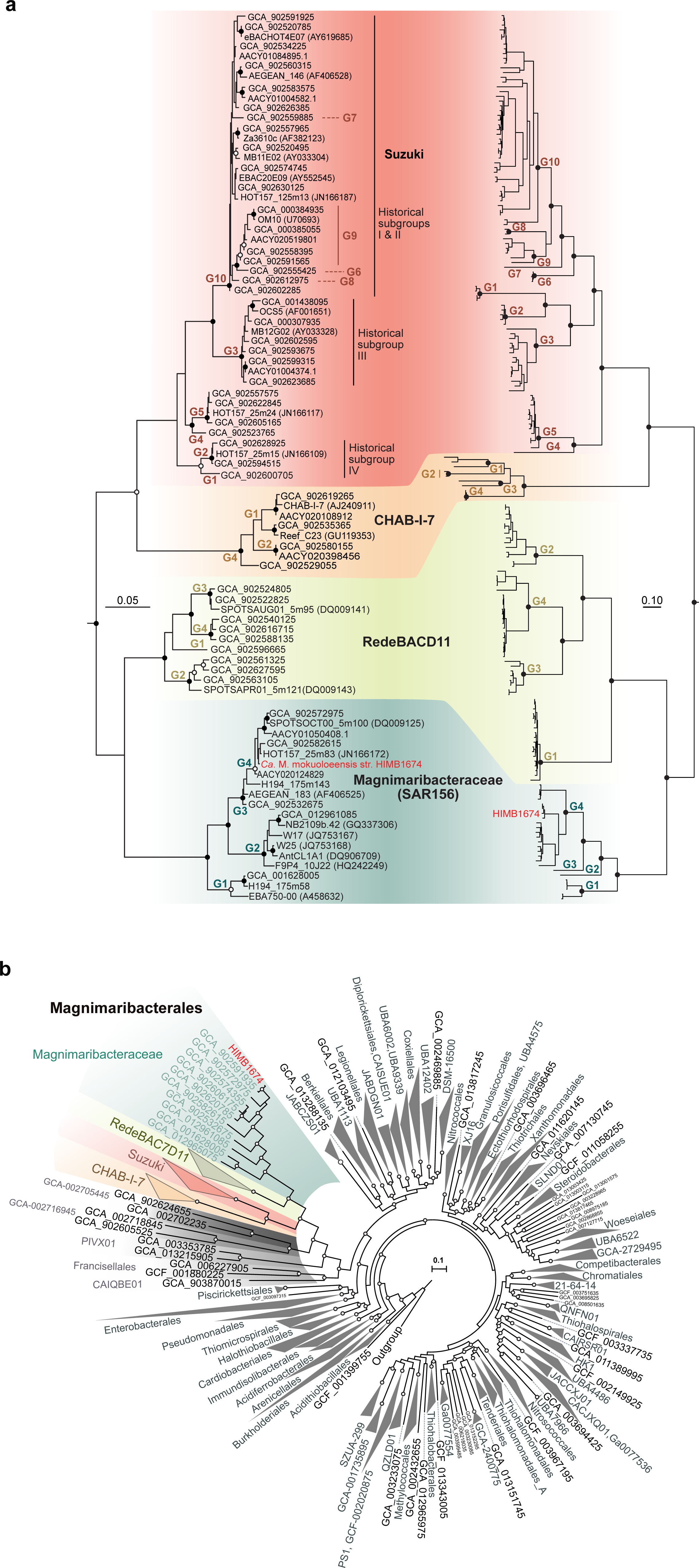
Phylogenetic placement of strain HIMB1674. a, Comparison between a 16S rRNA gene phylogeny (left) and a phylogenomic analysis from a concatenated alignment of 120 single copy core genes (right). *Ca.* Magnimaribacter mokuoloeensis str. HIMB1674 diverges at the root of SAR86 within the SAR156 subclade, now the family Magnimaribacteraceae. Historical subgroups CHAB-I-7 and RedeBAC7D1 are shown, along with historical subgroups I-IV that together make up the family-level Suzuki lineage. “G” denotes genus-level groupings defined by phylogenomics and RED values. Solid circles indicate bootstrap values greater than 95, while open circles indicate values over 80. The 16S rRNA gene phylogeny was constructed from rRNA genes extracted from the expanded SAR86 genomes dataset with historical sequences included for reference. Betaproteobacteria 16S rRNA gene sequences were used as an outgroup. The phylogenomic analysis included members of the gammaproteobacterial orders Burkholderiales and Pseudomonadales as an outgroup (GCF_000305785.2, GCF_003752585.1, GCF_003574215.1, GCF_006980785.1). For phylogenomic analysis, support value circles were removed from below genus nodes to maintain clarity on branching patterns. **b,** Phylogenetic placement of the SAR86 order Magnimaribacterales within the Gammaproteobacteria. Genomes from 416 family-level lineages of Proteobacteria from GTDB release 202 and 78 genomes from the Magnimaribacterales were included. The phylogeny is rooted at the internal node distinguishing the Gammaproteobacteria from the other three classes Alphaproteobacteria, Magnetococcia, and Zetaproteobacteria. Circles indicate ultrafast bootstrap support values ≥95%, from 1000 replicates.

**Figure 2.**
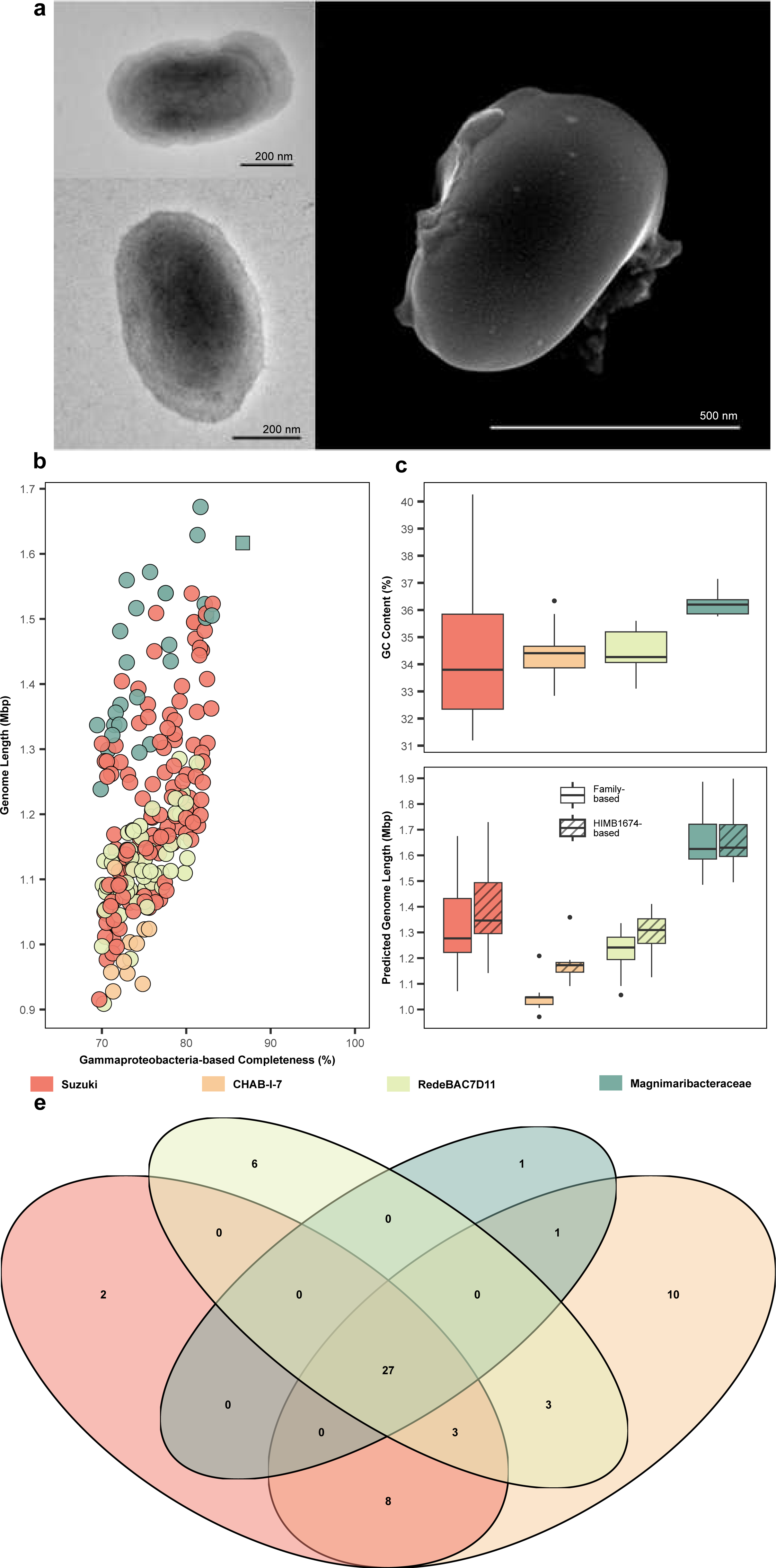
Characteristics of *Ca.* M. mokuoloeensis str. HIMB1674 and high quality SAR86 environmental genomes. a, TEM and SEM images reveal uniform small coccobacilli in a growing monoculture of strain HIMB1674. **b,** Distribution of genome completeness based on the GTDB Gammaproteobacteria marker gene set compared with absolute genome length. HIMB1674 is indicated by a square. **c,** The different families within the Magnimaribacterales harbor distinct %GC values. **d,** The different families within the Magnimaribacterales also harbor distinct predicted genome lengths, calculated with the marker genes present in the complete HIMB1674 isolate genome as well as marker genes present within the high quality genomes of each family. **e,** Venn diagram showing the overlap of missing marker genes from the Gammaproteobacteria marker gene set of CheckM. The SAR86 expanded genome dataset (n=185) used in panels b-e includes 25 Magnimaribacteraceae, 57 RedeBAC7D11, 10 CHAB-I-7, and 93 Suzuki genomes.

### A reduced marker gene set differentiates SAR86 from other Gammaproteobacteria

Although the genome of *Ca.* M. mokuoloeensis is closed, it lacks 28 of 280 genes in a widely used marker set for Gammaproteobacteria^26^ (Table 1), yielding a completion value of only 86.69%. Nearly all (27 of 28) of the missing genes are also systematically absent from a dataset of 732 putative environmental SAR86 genomes (Extended Data Table 1). A conservative correction of 28 missing genes that corresponds to the set absent from *Ca.* M. mokuoloeensis led to an increase in the average completion of the initial dataset of 705 putative environmental SAR86 genomes from just under 60% to 68% (Extended Data Table 2).

### SAR86 is a diverse lineage that branches deeply within the Gammaproteobacteria

From the initial dataset of 732 genomes, 224 were retained for downstream analyses following quality control (>80% complete and <5% contamination) using the modified marker gene set of 252 genes (Passed Quality Check in Extended Data Table 2). Together with the closed *Ca.* M. mokuoloeensis genome, this was further reduced to a set of 78 genomes that represent >93% ANI genome clusters (SAR86_species_rep in Extended Data Table 2), that we functionally refer to as species clusters.

A Proteobacteria-wide phylogenomic analysis constructed with a site-specific frequency model that included 771 reference genomes and the 78 genomes of the SAR86 species dataset revealed that nearly all of the putative SAR86 genomes grouped within a single monophyletic clade that includes HIMB1674 (Fig. 1b), with the exception of three genomes originating from the SAR86 ‘marine subsurface clade’ described by Zhou and colleagues^23^ that group within the bacterial order Pseudomonadales, where they are closely related to the marine gammaproteobacterial lineage OM182^8^ (data not shown). The SAR86 lineage is a deep, order-level branch within the Gammaproteobacteria, where it shares a common evolutionary origin with two lineages composed exclusively of metagenome assembled genomes (MAGs) generated from marine metagenomes, labeled GCA-002705445 and PIVX01 in the Genome Taxonomy Database (GTDB) release 202 (Fig. 1b). These three lineages of marine origin together share a common ancestry with the GTDB order CAIQBE01 that consists of two MAGs from freshwater lakes, and the order Francisellales, which includes a variety of cultured and uncultured microorganisms including representatives from freshwater and marine systems. We have named the SAR86 order-level branch of Gammaproteobacteria the Magnimaribacterales.

Using the SAR86 species dataset again (without the Pseudomonadales related genomes), taxonomic ranks assigned from relative evolutionary distance (RED) values yielded four families and 22 genera of Magnimaribacterales bacteria (Extended Data Fig. 1), consistent with the four families and 18 genera recognized in the GTDB. Cross-referencing with the 16S rRNA gene-based phylogeny revealed that 3 of the 4 family-level lineages corresponded to previously defined SAR86 subclades including RedeBAC7D11^27,28^, which has also been referred to as SPOTS^15^ (AG-339-G14 in GTDB), CHAB-I-7^27,29^ (TMED112 in GTDB) and SAR156^2,30^ (SAR86 in GTDB), which contains *Ca.* M. mokuoloeensis (Fig. 1). The fourth family-level lineage (D2472 in GTDB) consisted of the previously identified SAR86 subclades I, II, III, and IV^27,31^. In order to provide a meaningful label for this family-level lineage we use “Suzuki” after Marcelino T. Suzuki, in recognition of his influential research characterizing the diverse lineages of the SAR86 clade^27,31^. We also retain the original labels of RedeBAC7D11 and CHAB-I-7 for these families for consistency with previous literature, and name the SAR156 family-level lineage the Magnimaribacteraceae.

Phylogenomic analyses revealed that the family-level lineages Suzuki and CHAB-I-7 share a common evolutionary origin, as do the Magnimaribacteraceae and RedeBAC7D11 (Fig. 1). While the Magnimaribacterales appear to possess a rapid evolutionary clock relative to most other Gammaproteobacteria, this is particularly true for the CHAB-I-7 family (Fig. 1). Based on the geographic origin of genomes, the four families are widely distributed across the global ocean, but are mostly limited vertically to the ocean surface layer (Extended Data Table 2). The Magnimaribacteraceae are an exception; genomes from two Magnimaribacteraceae genera (G1 and G2) originate exclusively from below the photic zone, while the other two Magnimaribacteraceae genera (G3 and HIMB1674-containing Magnimaribacter) are affiliated with the surface ocean (Extended Data Table 2). This unique aspect of the Magnimaribacteraceae is corroborated by 16S rRNA gene analysis that revealed several lineages that originate from below the photic zone and in the deep ocean (Extended Data Fig. 2).

### Small genomes and low %GC typify SAR86 bacteria

An assessment of genome completeness based on the GTDB Gammaproteobacteria conserved gene set revealed completion values of all environmental genomes, barring those that belonged to the Pseudomonadales order, to be less than the 86.69% of HIMB1674 (Fig 2b). For this reason, we calculated modified completion estimates based off of the marker gene set adapted from the *Ca.* M. mokuoloeensis genome. Using this HIMB1674 conserved gene marker set, genomes within the four Magnimaribacterales families averaged 1.18 to 1.66 Mbp (Fig. 2d, Table 1, Extended Data Table 2). However, an analysis of the marker gene set across all of the major families within the Magnimaribacterales revealed that the likely number of missing marker genes ranged from 29 to 52 and was conserved at the family level (Table 1, Extended Data Table 1). When applying marker gene sets specific to each family, the average genome size decreased to 1.05 Mbp for CHAB-I-7 (Fig. 2d). The GC content of Magnimaribacterales genomes range from 31 to 40% (Extended Data Table 2), with some families and genera possessing distinct %GC (Fig. 2c, Extended Data Fig. 3).

### β-oxidation is central to the organic carbon metabolism of the Magnimaribacterales

Genes encoding enzymes that carry out the initiation and four steps of the β-oxidation pathway for fatty acid degradation are present in some of the highest copy numbers within the *Ca.* M. mokuoloeensis genome, and are also widespread across the Magnimaribacterales (Fig. 3, Extended Data Table 3). However, the CHAB-I-7 family lacked a 3-hydroxyacyl-CoA dehydrogenase gene required to fulfill step 3 (*fadB*), and generally did not show elevated copy numbers for this pathway. *Ca.* M. mokuoloeensis also encodes the ability to degrade the fatty acid-like side chains of steroidal compounds, including genes with homology to 3-oxocholest-4-en-26-oate-CoA hydrogenase (*fadE26,27*) and 3-oxochol-4-en-24-oyl-CoA hydrogenase (*fadE34*), that are also present in a majority of genera from the families Magnimaribacteraceae, Suzuki, and RedeBAC7D11 (Fig. 3, Extended Data Table 3). Similarly, *Ca.* M. mokuoloeensis contains putative Δ3,Δ2-enoyl CoA isomerases (ECI1_2) that are also present in a majority of Magnimaribacteraceae, Suzuki, and RedeBAC7D11 genomes, suggesting that *Ca.* M. mokuoloeensis and these three families share the ability to metabolize unsaturated fatty acids. This expands the variety of substrates that these lineages may degrade via β-oxidation. Magnimaribacterales genomes appear able to also obtain fatty acids from lipids through putative phospholipases found across all four families. A limited number of genomes also encode putative quorum quenching genes such as those for acyl-homoserine-lactone acylases (*quiP*) that also serve to provide another source of fatty acids. Finally, genes encoding putative phytanoyl-CoA dioxygenases (*phyH*) were identified within *Ca.* M. mokuoloeensis as well as other Magnimaribacteraceae, Suzuki, and RedeBAC7D11 genomes, revealing the potential ability to degrade the phytol sidechain of chlorophyll or similar compounds via α-oxidation (Fig. 3, Extended Data Table 3).

**Figure 3.**
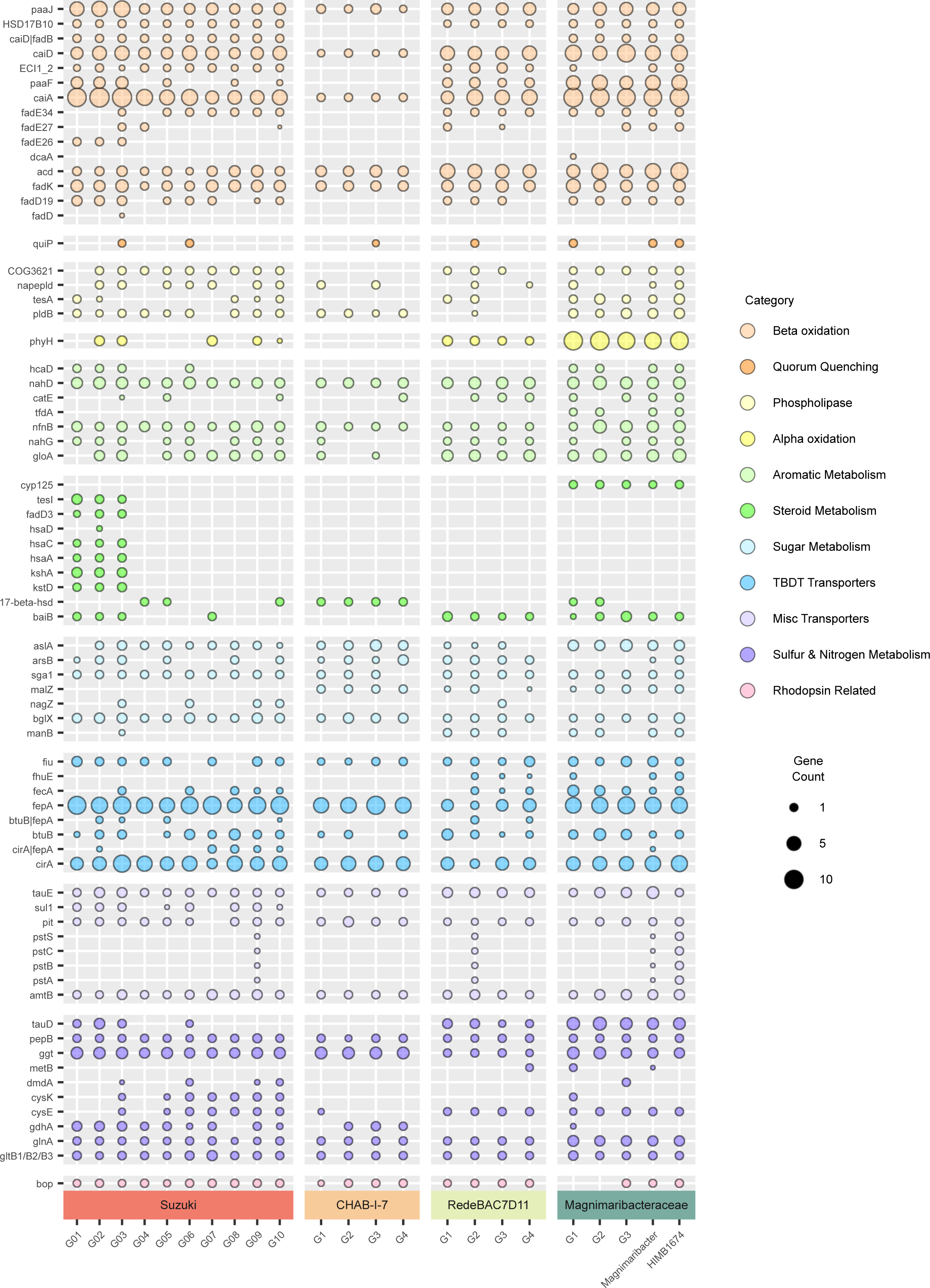
Distribution of key metabolic features across the Magnimaribacterales. The size of each circle indicates the average abundance of each gene within a genus. A comprehensive table of this data is provided in Extended Data Table 3.

### Aromatic and steroidal core ring cleavage is widespread across the Magnimaribacterales

In addition to its fatty acid side chain, a diverse set of genes encoding enzymes involved in the catabolism of the core rings of steroid-like compounds were identified across the Magnimaribacterales (Fig. 3, Extended Data Table 3). These include genes within all CHAB-I-7 and several Suzuki genera that encode putative 3(or 17)beta-hydroxysteroid dehydrogenases (17-beta-hsd) known to be involved in the degradation of sterols. In contrast, a majority of genomes from the RedeBAC7D11 and Magnimaribacteraceae families contain genes bearing homology to bile acid-coenzyme A ligases (*baiB*) that are known to play an important role in the degradation of bile acids^32^, which also possess a steroidal ring structure. Genes bearing homology to cholest-4-en-3-one 26-monooxygenases (*cyp125*), which participate in cholesterol ring degradation, were also identified in a majority of Magnimaribacteraceae genomes. Multiple sublineages within the Suzuki family contain genes encoding almost all the enzymes required to degrade androstenedione-like compounds into 3-[(3aS,4S,7aS)-7a-methyl-1,5-dioxo-octahydro-1H-inden-4-yl]propanoate and 2-hydroxy-cis-hex-2,4-dienoate in a majority of their genomes. 2-hydroxy-cis-hex-2,4-dienoate can be further degraded through the catechol meta-cleavage pathway (described below). Genes for this particular pathway were absent from the other genera-level lineages within Suzuki, as well as all other families of Magnimaribacterales.

In addition to steroidal compounds, the capacity to degrade aromatic compounds is likely widespread across the Magnimaribacterales (Fig. 3, Extended Data Table 3), and genomic evidence indicates that simple polycyclic aromatic hydrocarbons may be used as a substrate. Genes that encode putative 2-hydroxychromene-2-carboxylate isomerases (*nahD*), which are involved in the degradation of naphthalene to salicylate, are widespread across the Magnimaribacterales. In addition, genes encoding putative salicylate hydroxylases (*nahG*) that convert salicylate into catechol are present in a majority of Suzuki, RedeBAC7D11, and Magnimaribacteraceae genera. Genes encoding catechol 2,3 dioxygenases (*catE/gloA*), crucial to the meta-cleavage pathway of catechol degradation in order to complete aromatic ring cleavage, are a near-universal feature of Magnimaribacterales genomes and usually present in multiple copies. In addition to the widespread genes described above, the Magnimaribacteraceae contained genes encoding enzymes involved in aromatic compound degradation that were less widely distributed. This includes genes related to putative 3-phenylpropionate/trans-cinnamate dioxygenases (*hcaD*) involved in the metabolism of trans-cinnamate, and genes related to 2,4-dichlorophenoxyacetate dioxygenases (*tfdA*) that are linked to the degradation of 2,4-dichlorophenoxyacetic acid^33^. The potential for Magnimaribacterales bacteria to metabolize nitroaromatic substrates is supported by the presence of genes encoding putative nitroreductases (*nfnB*) that may degrade a broad range of nitroaromatic compounds, resulting in the release of nitrogen compounds such as ammonium (NH_4_) and potentially leading to the further degradation of the remaining aromatic end product via the gene complements described above^34^.

Sugar metabolism enzymes involved in the degradation of oligo- and polysaccharides were also encoded by Magnimaribacterales genomes (Fig. 3, Extended Data Table 3). While a majority of Magnimaribacterales encode glucoamylases (*sga1*) and beta-glucosidases (*bglX*), other enzymes are restricted in distribution. For example, genes encoding putative endo-1,4-beta-mannosidases (*manB*) are found across a majority of RedeBAC7D11 and Magnimaribacteraceae genomes but are largely absent from the other two Magnimaribacterales families. Genes encoding putative alpha-glucosidases (*malZ*) were present in the CHAB-I-7, RedeBAC7D11, and Magnimaribacteraceae families, but missing from the Suzuki family. Conversely, the Suzuki family was the only family-level lineage that contained genes encoding putative beta-N-acetylhexosaminidases (*nagZ*) across multiple genera. While the genomes of a single genus (G3) within the RedeBAC7D11 family also contained genes encoding putative beta-N-acetylhexosaminidases, they were not present within genomes of CHAB-I-7 and the Magnimaribacteraceae. All families within the Magnimaribacterales were also found to contain putative arylsulfatases (*aslA/arsB*) that are known to be involved in the desulphation of sulfoglycolipids.

Consistent with the genome of *Ca.* M. mokuoloeensis, all families within the Magnimaribacterales harbor the components necessary for a functional TCA cycle, glyoxylate shunt, and a complete 2-methylcitrate cycle (Fig. 3, Extended Data Table 3), except genomes of the CHAB-I-7 family which lack a gene encoding a methylisocitrate lyase (*prpB*). *Ca.* M. mokuoloeensis and other members of the Magnimaribacterales encode a complete Embden-Meyerhof-Parnas (EMP) glycolytic pathway, except that roughly half of RedeBAC7D11 and Magnimaribacteraceae genomes are missing a gene encoding glucose-6-phosphate isomerase (*pgi*) (Fig. 3, Extended Data Table 3).

### TonB-dependent transporters are prevalent within the Magnimaribacterales transporter pool

Genes encoding TonB-dependent transporters (TBDTs) were abundant across all families of the Magnimaribacterales (Fig. 3, Extended Data Table 3). Of these, two were particularly abundant: *cirA*, an iron-catecholate receptor, and *fepA*, a transporter for ferrienterochelin, the iron complex of the siderophore enterobactin. *FepA* has also been shown to transport vitamin B12 and some colicins. Other widespread TBDTs included *fiu*, a receptor for the iron siderophore ferrichrome A and monomeric catechols, and *fecA*, a receptor for ferric citrate that was particularly prevalent across RedeBAC7D11 and the Magnimaribacteraceae. The TBDT for vitamin B12, *btuB*, was also found throughout the Magnimaribacterales.

Outside of TBDTs, other prevalent transporters also provided clues regarding potential organic substrates used by Magnimaribacterales cells for growth. While several widespread transporters were annotated as sugar transporters, a close inspection revealed other potential substrates for transport that included cyclic compounds such as shikimate, and lipids such as sphingosine-1-phosphate (Fig. 3, Extended Data Table 3).

Genes encoding putative ammonia (ammonia channel protein *amtB*) and inorganic phosphate transporters (*pit*) were core features of Magnimaribacterales genomes (Fig. 3, Extended Data Table 3). The PsT system of high-affinity phosphate transporters was also present among certain RedeBAC7D11 and the *Ca.* M. mokuoloeensis genome.

### Ammonia and organic sulfur are likely sources of cellular nitrogen and sulfur across the Magnimaribacterales

Consistent with the presence of the ammonia channel protein (*amtB*) gene, mechanisms for nitrogen assimilation via ammonia were identified across the Magnimaribacterales. One pathway included putative glutamate synthase (*gltB1/B2/B3*) and glutamine synthetase (*glnA*) genes, which support the GS-GOGAT pathway (Fig. 3). The second mechanism, via putative glutamate dehydrogenases (*gdhA*), was almost exclusive to Suzuki and CHAB-I-7 (Fig. 3, Extended Data Table 3).

While no support for sulfate incorporation was found within Magnimaribacterales genomes outside of a putative sulfate permease across the Suzuki family, various alternative potential sources to fulfill cellular sulfur requirements were encoded by SAR86 genomes (Fig. 3, Extended Data Table 3). All four families within the Magnimaribacterales possessed putative gamma-glutamyltranspeptidases (*ggt*) and leucyl aminopeptidases (*pepB*) for converting glutathione into cysteine. In addition, almost all RedeBAC7D11 and Magnimaribacteraceae genomes, as well as a subset of Suzuki, possessed putative taurine dioxygenases (*tauD*) that may convert taurine into sulfite and other byproducts. However, Magnimaribacterales genomes lacked a recognizable sulfite reductase (*dsrA*, *cysJ*, *asrA*, *sir*) needed to convert sulfite into sulfide, and cysteine synthase (*cysK*) was largely limited to genomes from Suzuki and Magnimaribacteraceae G1. This suggests that the Magnimaribacterales are either unable to utilize the sulfite produced from taurine, utilize an unrecognized pathway for its utilization, or the putative taurine dioxygenase serves a different function. While cystathionine gamma-synthases (*metB*) would provide such an alternative method to incorporate sulfite, putative homologs were only found in genomes from RedeBAC7D11 and Magnimaribacteraceae, and not present across the Suzuki and CHAB-I-7 families. Finally, genes encoding putative dimethylsulfoniopropionate demethylases (*dmdA*) may confer the ability to degrade DMSP within a subset of genera primarily from the Suzuki family, providing another sulfur source for this limited number of genomes.

### Magnimaribacterales bacteria are primarily associated with surface seawater and make up ∼2-6% of the global ocean microbial community

Across a global survey of metagenomes spanning the upper 300 m of the water column, read recruitment revealed that the relative abundance of the Magnimaribacterales peaked in the surface ocean above 100 m and decreased with depth (Fig. 4a). However, the Magnimaribacteraceae were unique in that the relative abundance of the family peaked at depth (Fig. 4a). The depth-specific distribution was unevenly distributed across the four genera of the Magnimaribacteraceae, with two genera peaking below the upper euphotic zone of 0-100 m (Fig. 4b). Overall, relative abundance was unevenly distributed across genera, particularly within Suzuki, where some genera were either consistently more abundant across the global ocean, or abundant only in particular regions (Fig. 4c-f). For example, several genera that were largely absent from all other oceanic regions peaked in abundance within metagenomes from the Mediterranean Sea.

**Figure 4.**
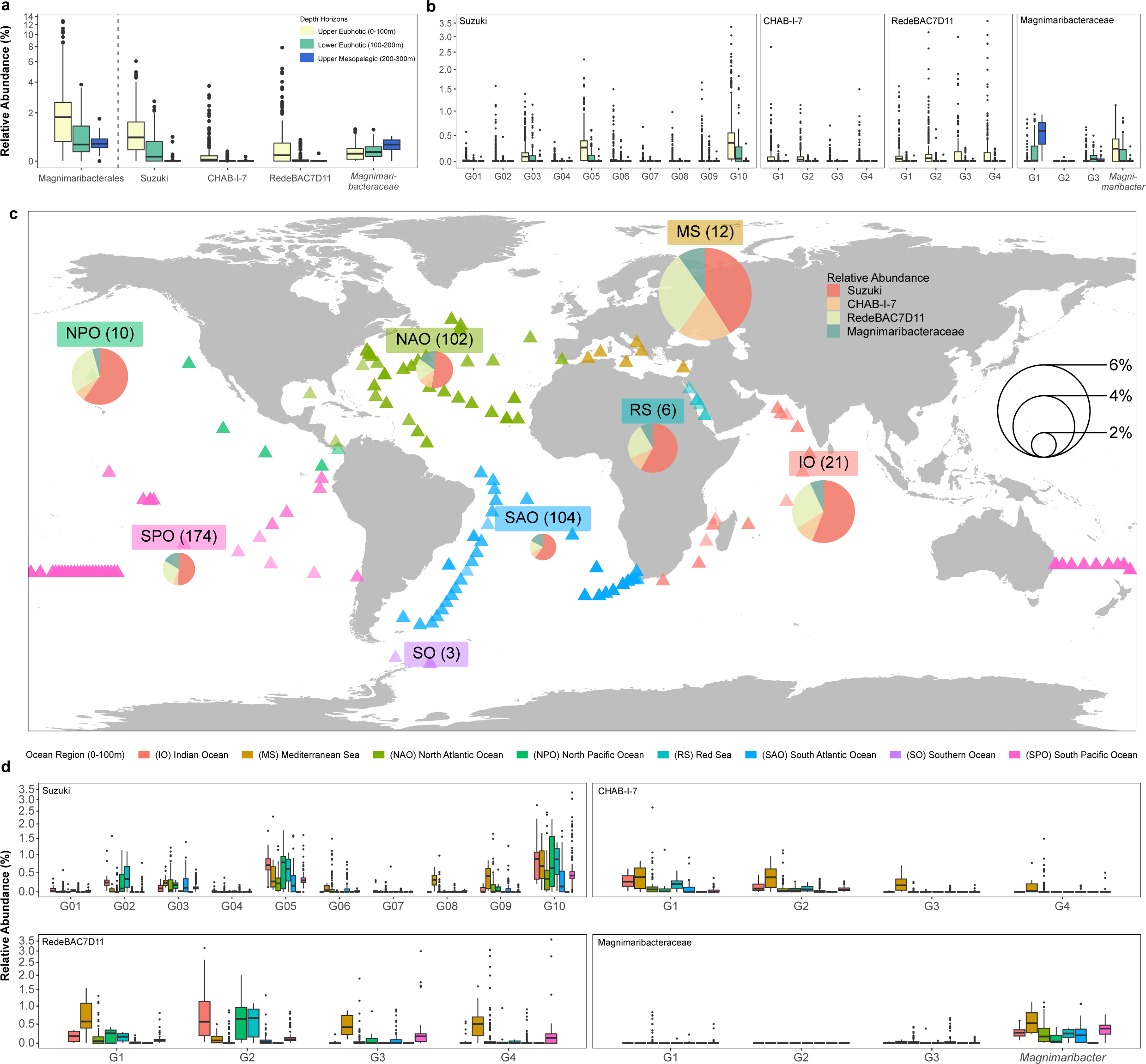
Relative abundance of Magnimaribacterales across the global ocean. a, The percent of reads recruited from the TARA (n=144) and GEOTRACES (n=462) metagenomes originating from depths above 300 m were mapped to genomes from across the Magnimaribacterales species dataset, revealing a clear preference for Magnimaribacterales within the uppermost layer (<100 m) of the water column. This was mirrored by the abundance of three of four families, with only the Magnimaribacteraceae peaking in abundance with depth. The legend indicates the depth horizon of the recruited metagenomes. **b,** Genera-level view of the read recruitment shown in panel a, highlighting differences in the distribution of genera and the prevalence of Magnimaribacteraceae G1 with depth. Relative abundance of Magnimaribacterales genera (**c-f)** and families (**g**) within the upper 100 m of the water column across the global ocean, using the read recruitment data shown in panel a. Oceanic regions are divided into the Indian Ocean (IO), Mediterranean Sea (MS), North Atlantic Ocean (NAO), North Pacific Ocean (NPO), Red Sea (RS), South Atlantic Ocean (SAO), Southern Ocean (SO), and South Pacific Ocean (SPO). Pie charts in panel g indicate the percent abundance of metagenomic reads recruited to the Magnimaribacterales species dataset. The numbers above each pie chart indicate the number of metagenomes utilized for the average relative abundance calculation from each region, while the size of the pie chart indicates the relative abundance of Magnimaribacterales in the particular oceanic region. Slices correspond to the relative abundance of each of the four families. Colored triangles indicate TARA Oceans and GEOTRACEs metagenome sample locations, with fill color corresponding to their associated ocean region.

Within the 0-100 m depth horizon, the relative abundance of Magnimaribacterales varied between different regions of the global ocean (Fig. 4g, Extended Data Table 4). While the Mediterranean Sea harbored the greatest average relative abundance of Magnimaribacterales bacteria (∼6%), other heavily sampled ocean regions such as the North Atlantic Ocean, South Atlantic Ocean, and South Pacific Ocean harbored fairly uniform Magnimaribacterales abundances of 2-3%. Magnimaribacterales were largely absent from the Southern Ocean. The average relative abundances of individual families of the Magnimaribacterales were stable across the global ocean, with the Suzuki family the most abundant across all oceanic regions.

Read recruitment to metagenomes that encompassed different size fractions did not reveal a clear trend that would indicate either extremely small cells, or those potentially associated with larger particles such as marine snow or ecto- or endosymbionts of larger cells (Extended Data Fig. 4).

### Description

In this study, we describe the first cultivar reported from the SAR86 lineage, HIMB1674, and propose the name “*Candidatus* Magnimaribacter mokuoloeensis” gen. nov., sp. nov., for this marine Gammaproteobacteria (Extended Data Table 5). The genus name Magnimari- is derived from the Latin words “magni” (here, great in quantity) and “mare” (sea). The species name mokuoloe-refers to the island (Moku o Loʻe) on which the laboratory is located where the strain was isolated. Accordingly, we also propose a new family name *Candidatus* Magnimaribacteraceae fam. nov., and order name *Candidatus* Magnimaribacterales ord. nov. to encompass all of the SAR86 lineage and harbor the genus and species defined here, *Ca*. Magnimaribacter mokuoloeensis.

## Discussion

Isolating the first strain of the SAR86 marine bacterial clade allowed us to assess the cellular, evolutionary, functional, and genomic characteristics of SAR86 within the context of a closed genomic template for the first time. A phylogenomic approach that accounted for the low %GC content of SAR86 genomes and included hundreds of newly available SAR86 single amplified genomes revealed that SAR86 forms an order-level lineage, the Magnimaribacterales, that branches deeply within the Gammaproteobacteria in a similar fashion to its SAR11 counterparts within the Alphaproteobacteria, the Pelagibacterales. Also consistent with other ubiquitous marine bacterial lineages such as the Pelagibacterales, Magnimaribacterales 16S rRNA genes form a broad clade composed of several sublineages widely distributed in the global ocean. However, historical 16S rRNA gene-based analyses have long revealed uncertainty with regard to affiliations within the Magnimaribacterales, and even what sublineages may or may not be part of this order. Our phylogenomic analyses clarify the presence of four robust family-level lineages within the Magnimaribacterales including the previously described CHAB-I-7 and RedeBAC7D11 families, the Suzuki family that includes previously described SAR86 subclades I through IV under its umbrella, and the SAR156 sublineage, which we describe here as the family Magnimaribacteraceae that contains *Ca.* Magnimaribacter mokuoloeensis str. HIMB1674. While historical studies have been inconsistent with their inclusion of the SAR156 sublineage within SAR86^15,27,35^, our phylogenomic analyses are consistent with the GTDB taxonomy and conclusively demonstrate that it forms a sister subclade to RedeBAC7D11 within the order Magnimaribacterales.

The capacity for β-oxidation was previously identified within a small set of incomplete Magnimaribacterales genomes^12,23^. Here we show that the β-oxidation pathway is not only found in all four families of the order (including within the genome of isolate *Ca.* M. mokuoloeensis), but multiple copies of the pathway are also present per genome across the Magnimaribacterales. We interpret the dedication of such a significant proportion of otherwise small genomes as an indication of the importance of this process to the core metabolism of Magnimaribacterales bacteria. Within individual genomes, the multiple gene copies at several steps of the β-oxidation pathway harbor considerable sequence variation. We hypothesize that these variants expand the variety of substrates available for degradation by this pathway. The presence of diverse lipases including phospholipase reveal the likely utilization of lipids as an organic substrate, with liberated fatty acids feeding into the β-oxidation pathway. Further, we infer it likely that the capacity to import into the cytoplasm and degrade a variety of lipid head group components is widely distributed across Magnimaribacterales genomes. This includes multiple genes encoding glycosidases that we predict localize within the periplasm and target glycolipids including lipopolysaccharides in order to cleave the oligo- or polysaccharide moiety, and genes encoding predicted arylsulfatase A enzymes capable of cleaving sulfate from sulfolipids prior to their further degradation into labile subunits. The sum of these reactions is the periplasmic degradation of complex lipids to generate relatively simple subunits that are subsequently available for import and degradation within the cytoplasm of Magnimaribacterales cells.

We were surprised to discover evidence that the Magnimaribacterales may also have the capacity to catabolize some components of steroidal compounds that, like lipids, are an important constituent of cell membranes. The ability to metabolize the side chain of the core four-ring steroid system and feed it through a series of β-oxidation reactions, similar to the degradation of lipid-derived fatty acids, appears to be a common trait of the Magnimaribacterales, with the exception of the CHAB-I-7 family. Perhaps more surprising is the presence of a small number of genus-level sublineages within the Suzuki family that possess the genetic potential to also fully degrade the core steroid rings and, in addition to these few genera, the broad distribution of several genes associated with the conversion or degradation of steroidal rings across the Magnimaribacterales. This suggests that the majority of Magnimaribacterales cells may also be able to catabolize at least a portion of the generally recalcitrant steroidal core.

Beyond steroid-like compounds, extensive genomic evidence indicates that cells of the Magnimaribacterales target other aromatic, heterocyclic, or polycyclic compounds for catabolism. The widespread presence of catechol 2,3-dioxygenases throughout the Magnimaribacterales indicates that the simple aromatic catechol is likely a significant substrate or intermediate for aromatic compound degradation, while the presence of ring-cleaving nitroreductases indicates that nitroaromatic compounds may also play an important role in Magnimaribacterales metabolism. Other lines of genomic evidence support our assertion that Magnimaribacterales cells specialize in the degradation of aromatic compounds. For example, the presence of 2-oxoglutarate ferredoxin oxidoreductases in an alternative TCA cycle, as found in the RedeBAC7D11 and Magnimaribacteraceae families, has been shown to facilitate the reduction of aromatic rings in anaerobic bacteria via an increase in production of reduced ferredoxin^36^. Glutathione S-transferases are known to play a role in the sequestration and transport of endogenous hydrophobic compounds, which includes steroids and other aromatic, heterocyclic, or polycyclic compounds. We hypothesize that the high copy number of genes encoding glutathione S-transferases across the Magnimaribacterales is a function of the important role they play in rendering some of these hydrophobic compounds more hydrophilic and thus accessible as substrates for catabolism by Magnimaribacterales cells. The high abundance of TonB-dependent transporters predicted to import iron-containing siderophores is also consistent with the high iron demand of enzymes involved in the predicted metabolisms of Magnimaribacterales, as iron is a common requirement of the ring-cleaving mono- and dioxygenases found in abundance across the order. Catechol serves as a subunit for several of the siderophores likely targeted by Magnimaribacterales TonB receptors, raising the intriguing possibility that these molecules are targeted for both iron and as a source of organic carbon.

Although the Magnimaribacterales is primarily a surface ocean-dwelling lineage of marine bacteria, the Magnimaribacteraceae family contains some sublineages that peak in relative abundance within the aphotic zone. Here we show that two of four genus-level sublineages within the Magnimaribacteraceae peak in abundance in the subsurface ocean, while the other two peak in the photic zone, including the genus *Ca.* Magnimaribacter. In support of this observation, genomes of the two genera of “deep” Magnimaribacteraceae systematically lacked a gene encoding the light-driven proton pump proteorhodopsin. This analysis thus provides a framework from which to investigate the depth-specific distribution of Magnimaribacterales genomes, and the genetic determinants of ecotypic differentiation with depth.

Across the global ocean, the relative abundance of Magnimaribacterales genomes in the upper 200 m of the water column ranged from an average of nearly 6% in the Mediterranean Sea to being absent from the Southern Ocean. However, SAR86 16S rRNA gene sequences have been found in seawater from both the Arctic and Antarctic^37^. Despite this variation in abundance, the relative proportions of the four families remained fairly stable across all ocean regions with Suzuki making up around half, RedeBAC7D11 making up around one fourth, and CHAB–I-7 and the Magnimaribacteraceae making up a little over 10% each of the global Magnimaribacterales community. This intriguing observation indicates that, while the proportion of Magnimaribacterales within a planktonic community may vary in relative abundance, the response of individual families is relatively stable and thus potentially predictable. Within each Magnimaribacterales family, a small subset of genera-level lineages were identified as the primary drivers of the relative abundance within each group. Two of ten and one (*Ca.* Magnimaribacter) of four genera were responsible for almost all of the read recruitment from most ocean regions within the Suzuki and Magnimaribacteraceae families, respectively. A unique exception to this was a select few genera-level lineages from Suzuki, CHAB-I-7, and RedeBAC7D11 that were almost completely absent from all other ocean regions except for the Mediterranean Sea. The presence of unique Mediterranean lineages is not exclusive to the Magnimaribacterales as it resembles previous observations of Pelagibacterales marine bacteria^38^. These Mediterranean specific sublineages also do not appear to directly compete with other closely related sublineages, as many of the most abundant sublineages found in the North Pacific and Indian Oceans are found at similar relative abundances in the Mediterranean. These results suggest that particular sublineages of the Magnimaribacterales possess characteristics adapted to niches only present in the Mediterranean.

The closed genome of *Ca.* M. mokuoloeensis lacked 10% of genes in the Gammaproteobacteria conserved marker gene set commonly used by the Genome Taxonomy Database to assess a variety of genome quality parameters, including estimated completion. This caused us to critically assess the presence of this marker set across the hundreds of environmental genomes investigated in this study, whereby we discovered that the Magnimaribacterales are systematically lacking nearly all (27 of 28) of the same makers, with additional marker sets consistently missing from each family. While the standard Gammaproteobacteria conserved marker gene set thus underestimates the completeness of Magnimaribacterales genomes and overestimates their predicted length, this is particularly true for the CHAB-I-7 lineage. This family systematically lacked 18% of the marker gene set that, when accounted for in completion estimates, results in genome length estimates of 1 Mbp. Intrigued by the unusually small genome size and fast evolutionary clock of this family relative to other Magnimaribacterales, we used read recruitment to different size fractions of marine plankton to tentatively conclude that cells of the CHAB-I-7 lineage are free-living bacteria and not unusually small in size or affiliated with larger cells. Coupled with apparent lesions in multiple metabolic pathways that are otherwise shared by other members of the Magnimaribacterales, these unique genomic characteristics make the CHAB-I-7 family a valuable target for understanding genome streamlining and the limits to genome reduction in presumably free-living microorganisms.

Magnimaribacterales and Pelagibacterales marine bacteria are two of the most abundant groups of heterotrophs inhabiting the global surface ocean where they share characteristic streamlined, low %GC genomes that harbor significant evolutionary divergence and concomitant genomic variation within their respective deep branches lineages of Proteobacteria. While genome-sequenced isolates have facilitated the dissection of Pelagibacterales metabolism for over two decades, our study represents the first report of a Magnimaribacterales isolate, its cellular characteristics, and its associated complete genome sequence.

One likely factor contributing to the cohabitation of Magnimaribacterales and Pelagibacterales cells is that they occupy different niche space. For heterotrophic bacteria in seawater, the diversity of compounds within the organic matter pool offers a plethora of opportunities for specialization and thus niche differentiation. While SAR11 has come to be known as a specialist in metabolizing labile, low-molecular-weight dissolved organic matter (DOM)^39^, we show that SAR86 instead targets a distinct and surprising fraction of the DOM including lipid and steroid components of cell membranes. The Magnimaribacterales are also likely capable of metabolizing other aromatic, polycyclic, and heterocyclic substrates. While the identity of these compounds is not yet known, hints gleaned from the gene annotations reveal catechol-based siderophones, chlorophyll, and nucleosides as potential targets. However, much of our metabolic inference is based on pathways described from human pathogens and microbes within other systems that are often distantly related to planktonic marine bacteria^40^, leaving open the opportunity describing specific substrates and the mechanisms by which Magnimaribacterales cells access and degrade this unique and frequently hydrophobic fraction of the global DOM pool. While we acknowledge that *Ca.* M. mokuoloeensis strain HIMB1674 is not yet sufficiently domesticated to the point where we are able to routinely assay for substrate utilization, the combination of complete genome sequence and an isolated strain provide the necessary tools needed to further this work.

## Methods

### Strain isolation, genome sequencing, and electron microscopy

The methods used to isolate strain *Ca.* M. mokuoloeensis strain HIMB1674 are described elsewhere^41^, with the following exceptions. A 4-liter seawater sample was collected on 26 July 2017 from station SB (N 21° 26.181’, W 157° 46.64196’) within Kāneʻohe Bay, Oʻahu, Hawaiʻi, and used as inoculum for a cultivation experiment, using the same sterile seawater growth medium described previously^41^. The raw seawater was diluted in medium to yield a cellular concentration of 2.5 cells per mL, aliquoted into six 96-well custom-made deep well Teflon plates, and incubated at 27°C in the dark. Between 3 to 8 weeks post-inoculation, the plates were screened for cellular growth as previously described^41^. Wells with growth of >10^4^ cells mL^-1^ were sub-cultured into 20 mL fresh medium, re-incubated, and subsequently screened again. Sub-samples of growing cultures were both cryopreserved and collected for DNA extraction, which was used to identify cultures via sequencing a small fragment of the 16S rRNA gene as previously described^41^. In order to more precisely determine the identity of *Ca.* M. mokuoloeensis str. HIMB1674, the full length 16S rRNA gene was also amplified, sequenced, and a phylogenetic trees created as described in the Supplementary Information.

The genome of *Ca.* M. mokuoloeensis strain HIMB1674 was sequenced (paired end, 2x151 bp) on a NextSeq 2000 (Illumina, San Diego, California, USA) from a library constructed using a modified Nextera XT library protocol^42^ and 7 ng of genomic DNA. After quality control using Trimmomatic v.0.36^43^, reads were assembled using metaSPAdes v.3.13.0^44^.

Scanning and transmission electron photomicrographs were prepared as described in the Supplementary Information.

### Environmental genomic data

Publicly available environmental genomes affiliated with the SAR86 lineage were identified within the Genome Taxonomy Database (GTDB)^45^ and PATRIC database^46^, and subsequently downloaded from Genbank. Within the GTDB, all genomes found under the bacterial order tag “o SAR86” were downloaded on 22 June 2020. The PATRIC database was searched for potential SAR86 genomes using the search term “SAR86” on 27 July 2020. Lastly, genomes characterized as SAR86 by the study of Zhou et al.^23^ were downloaded from Genbank. All genomes downloaded from Genbank utilized the bit program^47^.

Genomes were initially checked for quality and level of completion using the CheckM v1.1.2 taxonomy workflow and parameters for the class Gammaproteobacteria^26^. Of 280 marker genes, 252 were found within the closed *Ca.* M. mokuoloeensis strain HIMB1674 genome (Table 1, Extended Data Table 1). The 28 marker genes missing from HIMB1674 were subsequently excluded from a second iteration of the CheckM workflow to generate completion values based on HIMB1674, the only closed SAR86 genome standard. Subsequent analyses utilized completion estimates that were corrected by excluding Gammaproteobacteria marker genes missing from the HIMB1674 genome. Genomes that were >80% complete and contained <5% contamination based on this modified core marker gene set were retained for subsequent downstream analyses. The presence of Gammaproteobacteria marker genes were subsequently assessed across all genomes of the SAR86 lineage (Extended Data Table 1).

A recent publication reported on the assembly of complete SAR86 genomes from collections of closely related SAGs^22^. We attempted to verify the veracity of these assemblies and discovered them to be artificial chimeras resulting from extensive mis-assembly. We describe this further in the Supplementary Information. Thus, while some of the findings of this previous study are consistent with our current observations, we maintained reliance on the individual high-quality SAGs as they appear in their original publications so as to avoid artifacts arising from artificial, chimeric genomes.

### Phylogenomic analyses

Genomes were initially grouped into species clusters following the approach of Parks and colleagues^48^ using an average nucleotide identity (ANI) value of 93% as described in the with representative genomes from each species cluster referred to as the SAR86 species dataset (n=78 genomes, Extended Data Table 2). A Proteobacteria-wide phylogeny utilizing the SAR86 species dataset and reference genomes (Extended Data Table 6) was created in order to verify that these genomes form a monophyletic grouping to the exclusion of other Gammaproteobacteria, and to assess the evolutionary origins of the SAR86 lineage within the Gammaproteobacteria. Additional details are provided in the Supplementary Information.

To identify comparable taxonomic levels across sublineages of the Magnimaribacterales, the SAR86 species dataset was used but with the exclusion of genomes identified as not belonging to Magnimaribacterales. A relative evolutionary distance (RED) approach was then used to identify taxonomic levels within the SAR86 lineage^49^. Additional details are provided in the Supplementary Information.

In order to investigate evolutionary relationships within the SAR86 clade, an additional phylogenomic analysis using a broadened SAR86 dataset that included genomes sharing <99% ANI with their assigned species cluster representative, referred to as the SAR86 expanded dataset (n=188 genomes; Extended Data Table 2) was performed. The SAR86 expanded dataset and outgroup genomes were processed using the GTDB-Tk ‘identify’ and ‘align’ commands in the same manner as the Gammaproteobacteria-wide phylogeny. No reference genomes from GTDB were included during the alignment step. The alignment file was then used to infer a tree using Modelfinder within IQ-Tree^50^, which determined the best-fit model to be LG+F+R7 with 1,000 ultrafast bootstrap replicates. Genomes from members of the Gammaproteobacteria orders Burkholderiales and Pseudomonadales were used as an outgroup. Additional details of the phylogenomic analyses are provided in the Supplementary Information.

### Comparative genomics and read recruitment

The anvi’o pangenome workflow (v7.1)^51,52^ was used with the SAR86 expanded genome dataset to identify patterns in gene content. Metagenomic read recruitment was used to investigate the abundance of SAR86 genomes across two publicly available open-ocean sampling endeavors: TARA Oceans^53^ and GEOTRACES^54^ (Extended Data Table 7). The approach of Shaiber et al.^55^ was used to estimate the abundance of SAR86 genomes. Additional details are provided in the Supplementary Information.

## Acknowledgements

We thank Kawika Winter and Maria Chuvochina for their generous help in determining the name for the isolate described in this manuscript, A. Murat Eren for useful discussions, and the SeqCenter (formerly the Microbial Genome Sequencing Center) for sequencing the genome of HIMB1674. We thank Tina Weatherby of the University of Hawaiʻi at Mānoa Biological Electron Microscope Facility for their assistance and guidance with electron microscopy procedures. This research was supported by funding from the National Science Foundation to MSR (grants OCE-1538628 and OCE-2149128) and a Data Science Graduate Fellowship to OR (grant OAR-2118222 to G. Jacobs). This is SOEST contribution xxx and HIMB contribution xxx.

## Table Legends

**Table 1. Summary of the genomic characteristics of *Ca.* M. mokuoloeensis str. HIMB1674 and the four families of the order Magnimaribacterales.**

## Extended Data Table Legends

**Extended Data Table 1. Detection of the 280 gene markers included within the CheckM Gammaproteobacteria set across *Ca.* M. mokuoloeensis str. HIMB1674 and 731 putative environmental SAR86 genomes.** Markers included in the second iteration of the CheckM workflow are indicated in the HIMB1674_marker_set column.

**Extended Data Table 2. Characteristics of 732 putative environmental SAR86 genomes.** The SAR86_expanded column indicates genomes that were used in the SAR86 expanded dataset, while the SAR86_species_rep column indicates genomes used in the SAR86 species dataset. Columns labeled 0 - 5+ indicate the number of times each marker gene was identified in the genome. NA indicates data elements that were not included in publicly available metadata.

**Extended Data Table 3. Distribution of genes responsible for core metabolic pathways and functions across Magnimaribacterales genomes.** Columns ending with “Mean” indicate average copy number while those ending with “Proportion” indicate proportion of genomes with at least one copy. A column is present for each genus, as well as for the HIMB1674 and Pelagibacter str. HIMB083 genomes.

**Extended Data Table 4. Data underlying the estimated relative abundance of each Magnimaribacterales family shown in Figure 4g**.

**Extended Data Table 5. Protologue for *Candidatus* Magnimaribacter mokuoloeensis.**

**Extended Data Table 6. Genomes used to create the Proteobacteria-wide phylogenomic analysis shown in Fig. 1b**. Taxonomy classifications are from GTDB release 202. N/A indicates no available taxonomic classification.

**Extended Data Table 7. Metagenomes used to quantify the distribution of Magnimaribacterales genomes via read recruitment.** Oceanic regions correspond to those used in Figure 4. The columns read_pairs_raw and read_pairs_passed indicate the number of raw reads in each sample and the number of reads that passed quality control respectively.

## Extended Data Figure Legends

**Extended Data Figure 1.**
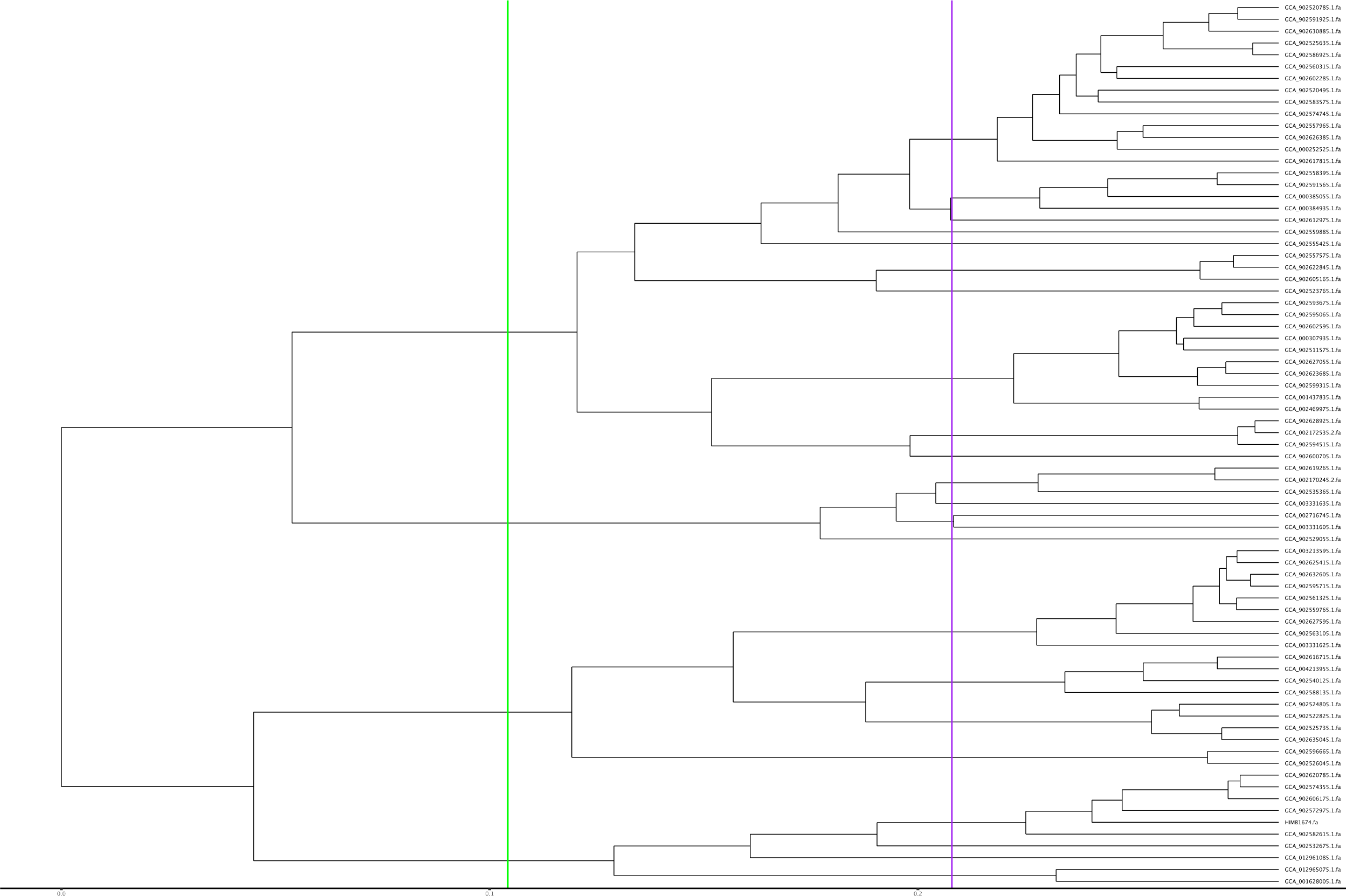
Relative Evolutionary Distance (RED)-scaled Magnimaribacterales phylogeny indicating family and genus designations. This phylum-wide analysis was performed using the Genome Taxonomy Database Toolkit (GTDB-Tk). Only the subtree for the Magnimaribacterales is shown. The green vertical line indicates the family-level RED value of 0.82, while the purple line indicates the genus-level RED value of 0.9237.

**Extended Data Figure 2.**
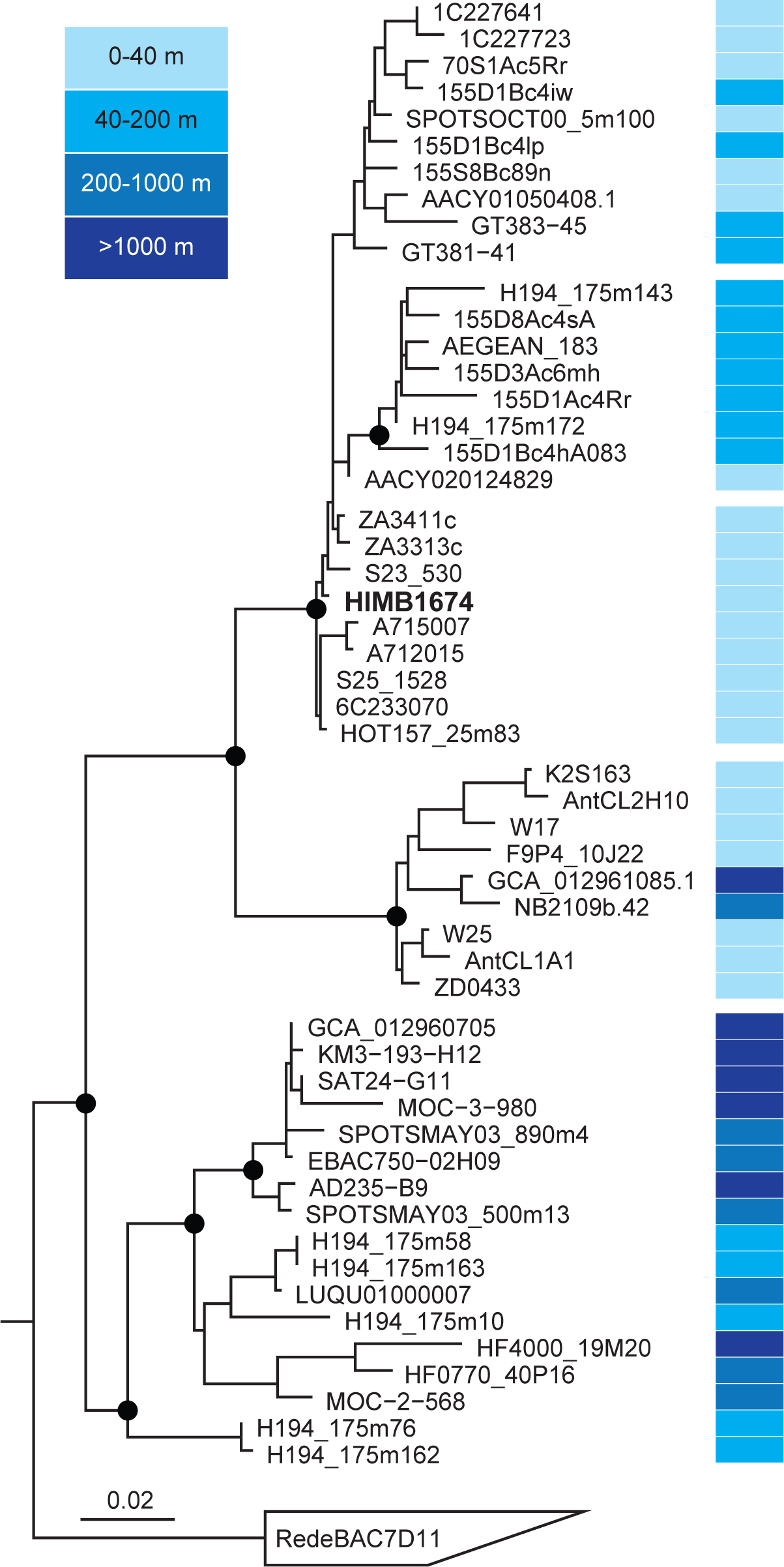
Ribosomal RNA-based phylogeny of the family Magnimaribacteraceae. Filled circles indicate major nodes with bootstrap support >90%, while colors indicate depth of sampling in the water column. The scale bar corresponds to 0.02 substitutions per nucleotide position. A selection of Betaproteobacteria were used as an outgroup.

**Extended Data Figure 3.**
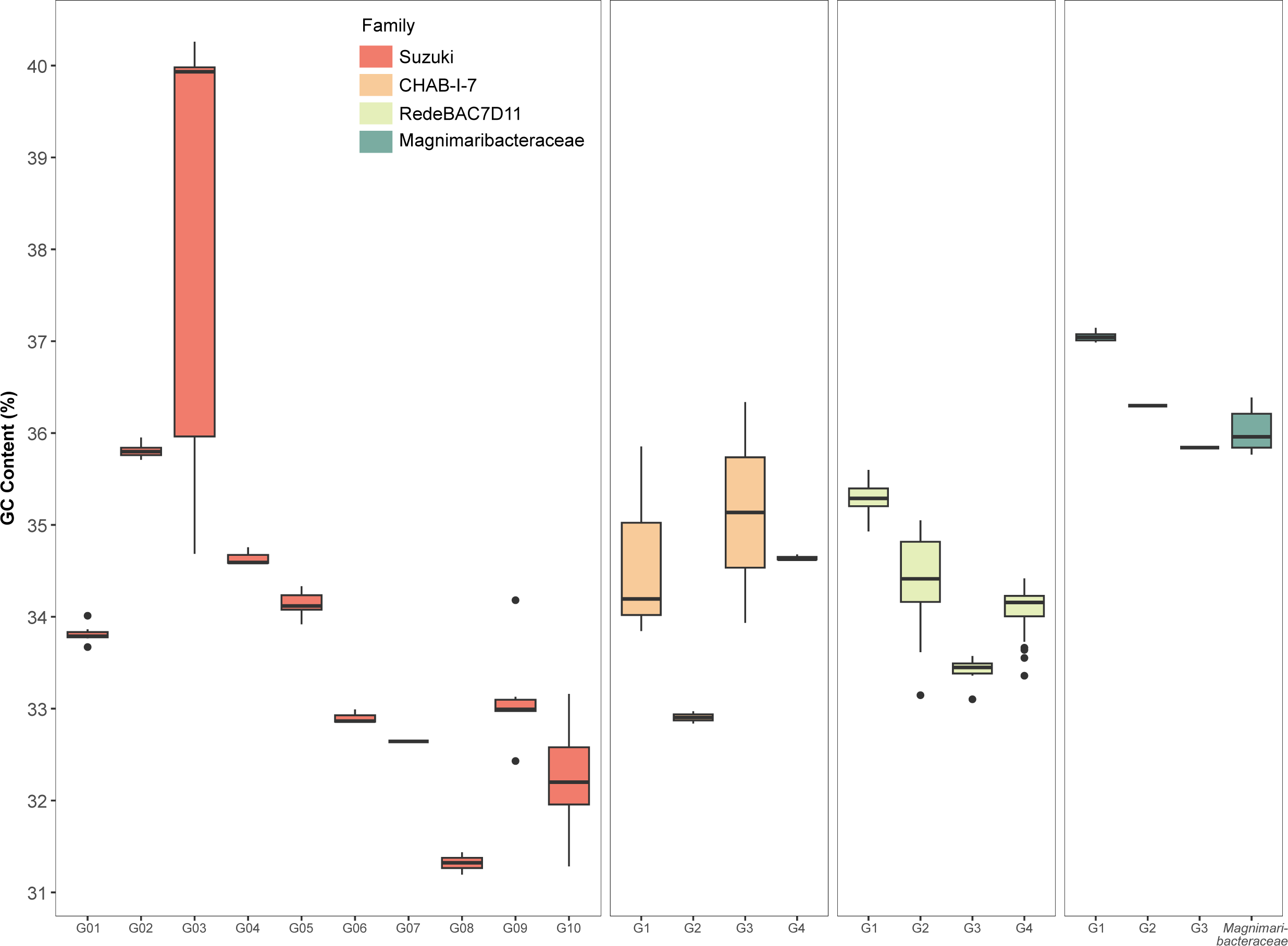
Distribution of GC content across genera of the four families of Magnimaribacterales. Color of box plots indicate the family that they belong to while the x axis indicates their genus.

**Extended Data Figure 4.**
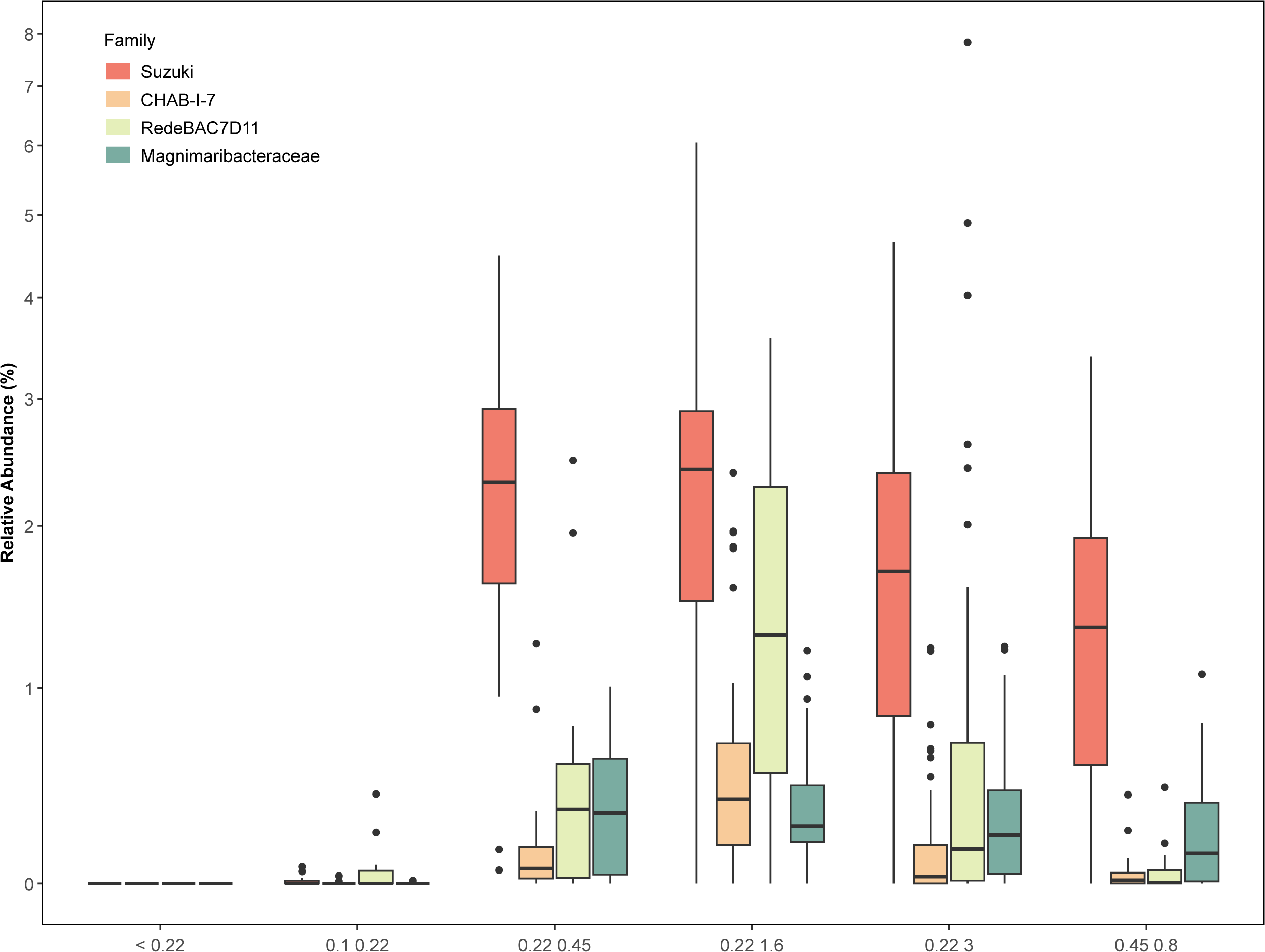
Relative abundance of metagenomic reads across the four Magnimaribacterales families within different size fractionated TARA Oceans metagenomes (n=204). Color of box plots indicate the family that each boxplot represents. The x axis indicates the different size fractionations that were present in the TARA oceans metagenomes used.

## Supplementary Information

### Electron microscopy

For preparation of specimens for transmission and scanning electron microscopy (TEM and SEM), 500 µL of cryopreserved HIMB1674 cells were defrosted at 20°C and fixed with glutaraldehyde (final concentration, 2.5%). The sample was concentrated 10-fold using a pre-rinsed Amicon Ultra-15, 30 kDa MWCO (Millipore Sigma) at 14,000 x g for 10 min, recovered per manufacturer instructions, and diluted to 80 µL in sterile seawater.

For TEM, 40 ul of sample was deposited onto a glow-discharged, formvar-coated mesh grid using an air-driven ultracentrifuge (Airfuge CLS, Beckman) as previously described^1^. Briefly, the grid was secured to the distal interior surface of the Airfuge rotor chamber (EM-90, Beckman) and samples were centrifuged for 24 min at 20 PSI. After centrifugation, grids were stained with 0.02 µm-filtered 2% uranyl acetate. Grids were examined with a Hitachi HT7700 transmission electron microscope with 100 kV accelerating voltage. High resolution images were taken using an AMT BioSprint16M-Active Vu 16 megapixel cooled 4896 x 3920 pixel CCD camera.

For SEM, fixed HIMB1674 cells were loaded onto 13 mm diameter, 0.1 µm pore-sized polycarbonate filters, washed twice with 0.1 M sodium cacodylate buffer with 0.44 M sucrose, postfixed with 1% OsO4 in 0.1 M sodium cacodylate, and finally dehydrated through a graded ethanol series (30%, 50%, 70%, 85%, 95%, 100%). Filters were submerged in 1:2 EtOH/HMDS (hexamethyldisilazane) for 40 min, then left in 100% HMDS overnight until dry. Filters were mounted on aluminum stubs with double-stick conductive carbon tape and coated with gold/palladium in a Hummer 6.2 sputter coater. Specimens were viewed and digital images were acquired with a Hitachi S-4800 Field Emission Scanning Electron Microscope at an accelerating voltage of 5 kV. TEM and SEM micrographs were analyzed using ImageJ software^2^. Cell biovolume was calculated using the equation for prolate spheriod cells^3^: *V = π / 6 ^·^ d^2·^ w*

### PCR amplification and phylogenetic analysis of the full length SSU rRNA gene

Approximately 35 picograms of genomic DNA from HIMB1674 was used as template for PCR amplification (Bio Rad C1000 Touch; Bio Rad, Hercules, CA, USA) using the primers 27FB^4^ and R1492^5^ that targeted the full length 16S rRNA gene. The 25 μL reaction volume contained 1 μL of genomic DNA template, 0.5 μL of each forward and reverse primer (final concentration of 0.2 µM), 10 μL of 5PRIME HotMasterMix (Quantabio, Beverly, MA, USA), and 13 μL of H_2_O. The reaction included an initial denaturing step of 2 min at 94°C, followed by 35 cycles of 30 s at 95°C, 1 min at 51°C, and 45 sec at 72°C, and a final extension of 12 min at 72°C. The resulting PCR product was cleaned (QIAquick PCR Purification Kit), Sanger sequenced in the forward and reverse directions, and assembled into a single contig.

The full length 16S rRNA gene from strain HIMB1674 was imported into the ARB software package^6^ along with those extracted from the environmental genomes of the SAR86 expanded dataset where available. The genes were aligned using SINA v.1.2.11^7^ to a curated database of marine 16S rRNA gene sequences based on Silva v.95^8^, and phylogenetic analyses were performed using the RAxML maximum likelihood method with the GTR model of nucleotide substitution under the gamma and invariable-models of rate heterogeneity^9^. Bootstrap analysis (1,000 replicates) was performed using the rapid bootstrap analysis algorithm of RAxML. A set of Betaproteobacteria 16S rRNA gene sequences from cultured isolates were used as an outgroup.

### Evaluation of “complete composite genomes” reported by Rodo-Garcia et al

We used multiple approaches in an attempt to verify that the “complete composite genomes" (CCGs) reported by Roda-Garcia and colleagues are circularized and single contig as reported^10^. First, we attempted to repeat the assembly of CCGs from their respective SAGs using the program Flye v2.9^11^ with the arguments ‘-genome size 1.3m -subassemblies’ as reported by Roda-Garcia and colleagues. Of the nine assemblies, three were circularized and single contig, one was a single open contig, and six contained multiple contigs (Supplementary Table 1). We also attempted to verify the validity of the nine original CCGs reported by Roda-Garcia et al.^10^ using a sequence mapping approach. First, we fragmented the original SAGs used to assemble each CCG into 10 kb fragments with 1 kb overlap, and 1 kb fragments with 100 bp overlap, using the tool cut_up_fasta.py from CONCOCT^12^. We then mapped the resulting 10 kb and 1 kb fragments to their respective CCG separately and in combination, and inspected the mapping manually. Numerous mis-assemblies were clearly evident across each of the nine CCGs, ranging from 6 in TMED112-A1 to 31 in D2474-A1-2 (Supplementary Table 1). Close inspection revealed that, at least in this case of this study, the Flye assembler had joined sequences erroneously. The Flye Github documentation also revealed that the ‘-subassemblies’ option is not as thoroughly tested as the primary workflows and was deprecated in v2.9. Given the convincing evidence that the CCGs of Roda-Garcia et al.^10^ are chimeric artifacts, we were compelled to use the native SAGs generated by Pachiadaki et al.^13^ in our analyses.

### Phylogenomic analyses

To create species clusters, the program fastANI v1.32^14^ was first used to calculate average nucleotide identity (ANI) between all quality-filtered Magnimaribacterales genomes. A histogram of ANI values identified a peak in ANI at 93-94% (Supplementary Fig. 1), and so an ANI value of 93% was used to group the Magnimaribacterales genomes into species clusters. In order to create species clusters, the genomes were sorted from maximum to minimum percent completeness. Then, starting with the most complete genome, a species cluster was created by identifying all other Magnimaribacterales genomes with >93% ANI relative to this representative. After removing the clustered genomes from the rank completion list, the genome with the next highest completion value was used in the same manner to create a species cluster. Through this iterative process, all quality-controlled genomes were grouped into individual species clusters that each possessed a representative genome of the highest completion.

In order to create the Proteobacteria-wide phylogeny with the Magnimaribacterales species dataset, marker genes for the Magnimaribacterales species dataset were first identified using the GTDB-Tk v1.7.0 ‘identify’ program^15^, which used Prodigal v2.6.3^16^ to call genes and HMMER v3.1b2^17^ to identify the 120 bacterial marker genes used by GTDB^18^. GTDB-Tk ‘align’ was then used to create a multiple sequence alignment of the Magnimaribacterales species dataset and GTDB-identified reference genomes for every species cluster within the GTDB phylum Proteobacteria using the 120 marker genes identified above. IQ-Tree v2.1.2 was used to perform a phylogenomic analysis using this alignment file with the LG4X+F model^19^. Using the large phylogeny as a guide, Treemmer v0.3^20^ was subsequently implemented to prune the tree to 1,000 genomes, with the conditions that at least two genomes from each family of *Proteobacteria* were retained (if available) and no Magnimaribacterales representative genomes were removed. The genomes retained in the pruned genome alignment were used as input for GTDB-Tk ‘identify’, GTDB-Tk ‘align’ (with the "–skip_gtdb_refs" flag), and IQ-Tree (LG4X+F model). This phylogeny was then used as the guide tree to calculate site frequency profiles for another phylogenomic analysis that employed a site-specific frequency model (model Poisson+UDM0064NONE+G4). This analysis used the UDM 64 component model with no transformations (UDM0064NONE) from the Homology-Derived Structures of Proteins (HSSP) database^21^, with IQ-Tree used to construct the phylogeny. This phylogeny was inspected to identify duplicate genomes, genomes labeled as “SAR86” but evolutionarily unaffiliated with the SAR86 lineage, and MAGs outside of Magnimaribacterales that possessed anomalously long terminal branches. After removal of these genomes, 849 remained (Table S3). The phylogeny produced using the UDM 64 site-specific frequency model was subsequently regenerated after removing the anomalous genomes, using 1,000 ultrafast bootstrap replicates.

To identify comparable taxonomic levels across sublineages of the Magnimaribacterales, the Magnimaribacterales species dataset was used. A relative evolutionary distance approach was then used to identify taxonomic levels within the SAR86 lineage. A bacterial domain-level phylogeny was created using the GTDB-Tk ‘de_novo_workflow’^15^ with the SAR86 expanded dataset and “p Chloroflexota’’ as the outgroup. This workflow identified marker genes in the input genomes and aligned them using the same methods previously described. The workflow then inferred a tree using FastTree v2.1.10 (model WAG+GAMMA)^22^, rooted the tree on the specified outgroup taxon (p Chloroflexota), and decorated the internal nodes using the GTDB taxonomy. This phylogeny was then used as the input for the scale_tree program in PhyloRank v0.1.11 (https://github.com/dparks1134/PhyloRank) to convert branch lengths into relative evolutionary distance (RED). RED values of 0.82 and 0.9237 were used to identify family and genus-level lineages, respectively. These values were based on the distribution of internal nodes within the Magnimaribacterales clade, their support values, and values used previously for other family and genus-level lineages^18^.

### Read recruitment

Metagenomic read recruitment was used to investigate the abundance of Magnimaribacterales genomes across two publicly available open-ocean sampling endeavors: TARA Oceans^23^ and GEOTRACES^24^. Within the TARA Oceans study, only metagenomes sequenced from the 0.22-3 μm size fraction were used. To prevent closely related genomes from competing for reads, only the Magnimaribacterales species dataset was used. Sequence reads were quality-filtered using ‘iu-filter-quality-minoche’ from the illumina-utils program v2.12^25^. Bowtie2 v2.4.2^26^ was used to map reads from the metagenomes to the genomes. SAMtools v1.11^27^ was used to sort and index SAM files into BAM files. To obtain detection estimates, a contig database was created from the concatenated genomes and used to profile each of the BAM files using ‘anvi-profile’ from the anvi’o program suite v7.1^28^. These profiles were then merged into a single profile database. Contigs in the merged profile were linked back to their associated genome using published methods^29,30^. The merged profile was then summarized using ‘anvi-summarize’ to obtain statistics on detection and abundance for the genomes in every metagenome.

The approach of Shaiber et al.^31^ was used to estimate the abundance of Magnimaribacterales genomes, calculated by taking the number of mapped reads divided by the total number of quality controlled reads found for each metagenome. These values were then divided by the size of each genome. This calculation was also performed for reads that did not map to any of the Magnimaribacterales genomes. For this unidentified fraction, an average genome size of 1.6 Mbp was used, calculated from the relative abundance and genome sizes found by Pachiadaki and colleagues^13^. Relative abundances were estimated by summing together the normalized abundances of the Magnimaribacterales genome and the unidentified fraction for each metagenome. The sum was then divided from each normalized abundance estimate to generate the relative abundance of the genome or the unidentified fraction in the metagenome. To correct for genomes that had low detection values (i.e. breadth of coverage), the relative abundance of any genomes that had a detection value below 50% was defaulted to 0, indicating that the genome was not present in the associated metagenome. This detection threshold value was based on an analysis previously performed on *Prochlorococcus* populations^30^. To calculate the relative abundance of family and genus-level lineages within the Magnimaribacterales, the relative abundance of each genome within each family and genus-level lineage was summed.

### Pangenomics

The anvi’o pangenome workflow (v7.1)^28,30^ was used with the Magnimaribacterales expanded genome dataset to identify patterns in gene content. A contig database was created for each Magnimaribacterales genome and annotated with clusters of orthologous genes (COGs)^32^ and Kyoto’s Encyclopedia of Genes and Genome^33^ using ‘anvi-run-ncbi-cogs’ and ‘anvi-run-kegg-kofams’, respectively. The module ‘anvi-run-hmms’ was used to identify a collection of bacterial single copy core genes, which identified open reading frames using Prodigal^16^ and identified them via HMMER^17^. Subsequently, a genome storage database was generated using the program ‘anvi-gen-genomes-storage’. The genome storage database was used as input for the pangenomic analysis using the command ‘anvi-pan-genome’ with the parameters "–minbit 0.5", "–mcl-inflation 2", and "–use-ncbi-blast". This command used BLAST^34^ to examine similarity between pairs of genes using the translated DNA sequence and the Markov Cluster algorithm^35^ to form homologous gene clusters.

## Supplementary Results

### The TCA and methylcitrate cycles are important components of central carbon metabolism in SAR86 bacteria

All Magnimaribacterales families contain the components necessary for a functional TCA cycle (Extended Data Table 3). While genomes of the Magnimaribacterales lacked a gene for citrate synthase, they contained a 2-methylcitrate gene that can fulfill the same function as citrate synthase while also playing a role in the 2-methylcitrate cycle. The Suzuki and CHAB-I-7 families both contained genes encoding the components of the 2-oxoglutarate dehydrogenase complex responsible for the conversion of 2-oxoglutarate to succinyl-CoA; however, these were largely absent in Magnimaribacteraceae and RedeBAC7D11 which instead possessed a 2-oxoacid oxidoreductase complex that likely fulfills this function (Fig. 3, Extended Data Table 3)^36^. General features of Magnimaribacterales genomes also include both components of the glyoxylate shunt as well as a complete 2-methylcitrate cycle, except for the CHAB-I-7 family which lacked a gene encoding a methylisocitrate lyase needed to complete 2-methylcitrate cycle. This suggests that CHAB-I-7 may lack the capacity to convert methylisocitrate to pyruvate and succinate.

While most components of the Embden-Meyerhof-Parnas (EMP) glycolytic pathway were present across the Magnimaribacterales, a gene encoding glucose-6-phosphate isomerase was absent from many RedeBAC7D11 and Magnimaribacteraceae genomes (Fig. 3, Extended Data Table 3). RedeBAC7D11 also lacked a pyruvate kinase, suggesting that this family may not have the capacity to convert phosphoenolpyruvate to pyruvate. The genes required for gluconeogenesis were present across all families of the Magnimaribacterales (Fig. 3, Extended Data Table 3). While most genomes from the Suzuki and CHAB-I-7 families contained genes encoding the components for a pyruvate dehydrogenase complex, RedeBAC7D11 and Magnimaribacteraceae lacked genes encoding the E1 and E2 components. Instead, these two families may utilize the same 2-oxoacid oxidoreductase complex used in the TCA cycle in order to convert pyruvate into acetyl-CoA (Fig. 3, Extended Data Table 3).

### Magnimaribacterales genomes harbor genes to produce bacteriorhodopsin but not all-*trans*-retinal

Genes encoding proteorhodopsin were ubiquitous across the Magnimaribacterales except for G1 and G2 of the Magnimaribacteraceae (Fig. 3, Extended Data Table 3). All genomes of the Magnimaribacterales lacked the ability to synthesize retinal de novo. While predicted all-trans-8’-apo-beta-carotenal 15,15’-oxygenases found across the Magnimaribacteraceae and within some genomes of the RedeBAC7D11 family suggests at least a portion of the Magnimaribacterales is capable of cleaving β-carotene to produce retinal, the presence of these functions in Magnimaribacterales genomes that lack the gene for proteorhodopsin raises the possibility that it serves an alternative function.

## Supplementary Figure Legends

**Supplementary Figure 1.**
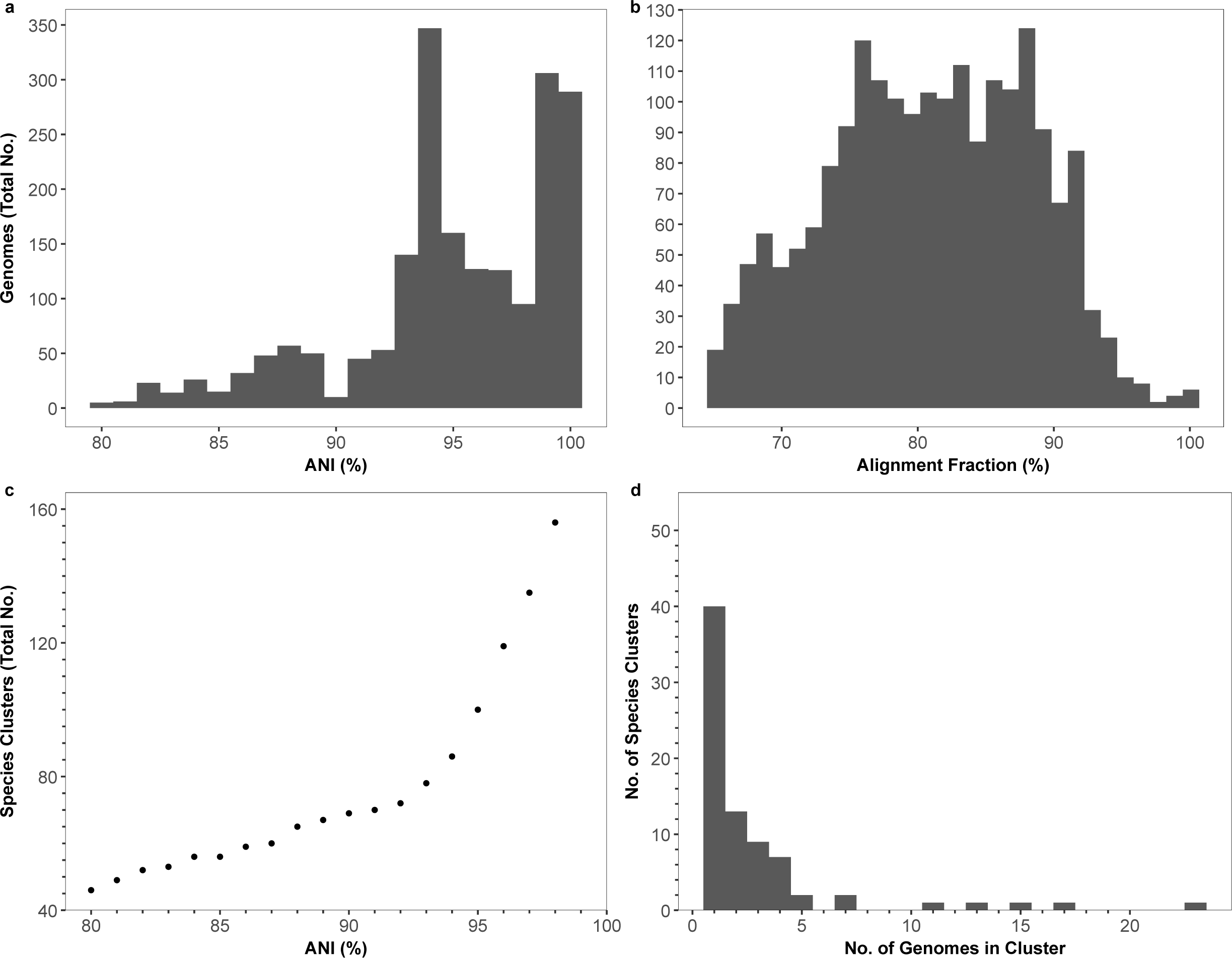
Characteristics of pair-wise comparisons between quality filtered Magnimaribacterales genomes (n=224). a, Histogram of average nucleotide identity (ANI) values (%) among all quality filtered SAR86 genomes, revealing a peak at 93-94%. **b,** Histogram of the alignment fraction (%) between all quality filtered SAR86 genomes. **c,** Number of species clusters created for different ANI values. **d,** Histogram of the distribution of genomes within each species cluster using ANI value of 93%.

## Supplementary Table Legends

**Supplementary Table 1. Summary of assembly errors in CCGs (complete composite genomes) reported by Roda-Garcia et al. (2023).**

